# Morphologically Constrained and Data Informed Cell Segmentation of Budding Yeast

**DOI:** 10.1101/105106

**Authors:** Elco Bakker, Peter S. Swain, Matthew M. Crane

## Abstract

**Motivation:** Although high-content image cytometry is becoming increasingly routine, processing the large amount of data acquired during time-lapse experiments remains a challenge. The majority of approaches for automated single-cell segmentation focus on flat, uniform fields of view covered with a single layer of cells. In the increasingly popular microfluidic devices that trap individual cells for long term imaging, these conditions are not met. Consequently, most segmentation techniques perform poorly. Incorporating information about the microfluidic features, media flow and morphology of the cells can substantially improve performance, though it may constrain the generalizability of software.

**Results:** Here we present DISCO (Data Informed Segmentation of Cell Objects), a framework for using the physical constraints imposed by microfluidic traps, the shape based morphological constraints of budding yeast and temporal information about cell growth and motion, to allow tracking and segmentation of cells in micrflouidic devices. Using manually curated data sets, we demonstrate substantial improvements in both tracking and segmentation for this approach when compared with existing software.

**Availability:** The MATLAB^®^ code for the algorithm and for measuring performance is available at https://github.com/pswain/segmentation-software. The test images and the curated ground truth results used for comparing the algorithms are available at http://swainlab.bio.ed.ac.uk/.

## Introduction

One of the primary methods through which information is acquired from biological samples is by optical imaging. Imaging by both transmitted and fluorescent methods is essential to the modern biological research laboratory, and the proliferation of innovative imaging techniques continues to increase its importance ([Meijering et al.(2016) Meijering, Carpenter, Peng, Hamprecht, and Olivo-Marin]). The automated application of these imaging methodologies, often in time-lapse microscopy experiments, has left biomedical researchers with a deluge of image data, and a common bottleneck to scientific analysis is the necessary segmentation into cells or regions of interest.

This challenge has been widely recognized for nearly fifty years, and has been the subject of intense research efforts ([Meijering et al.(2016)]). The most widely used and most generalizable methods rely on thresholding images into a foreground and background ([Kamentsky et al.(2011)Kamentsky, Jones, Fraser, Bray, Logan, Madden, Ljosa, Rueden, Eliceiri, and Carpenter]). Although useful, these methods have several problems that warrant the development of tools tailored to specific uses ([Sommer and Gerlich(2013)]). Most importantly, to achieve an acceptable accuracy a fluorescent marker is typically used to label part of the cell ([Schiegg et al.(2015)Schiegg, Hanslovsky, Haubold, Koethe, Hufnagel, and Hamprecht, Federici et al.(2012) Federici, Dupuy, Laplaze, Heisler, and Haseloff, Zhong et al.(2012)Zhong, Busetto, Fededa, Buhmann, and Gerlich, Kamentsky et al.(2011) Kamentsky, Jones, Fraser, Bray, Logan, Madden, Ljosa, Rueden, Eliceiri, and Carpenter, Conrad et al.(2011)Conrad, Wünsche, Tan, Bulkescher, Sieckmann, Verissimo, Edelstein, Walter, Liebel, Pepperkok, and Ellenberg, Held et al.(2010)Held, Schmitz, Fischer, Walter, Neumann, Olma, Peter, Ellenberg, and Gerlich, Pelet et al.(2012)Pelet, Dechant, Lee, van Drogen, and Peter]), which both increases the workload of constructing strains and occupies a fluorescent channel: limiting the amount of data that can be acquired. For example, a large proportion of the ORF-GFP library in budding yeast ([Huh et al.(2003)Huh, Falvo, Gerke, and Carroll]), over four thousand cell lines, were tagged with red fluorescent protein purely to facilitate automated segmentation ([Chong et al.(2015)Chong, Koh, Friesen, Duffy, Cox, Moses, Moffat, Boone, and Andrews]). There is thus an urgent need for a reliable segmentation method based on the readily obtainable bright field or DIC images that would prevent this additional work.

Image segmentation methods, and computer vision in general, have to balance trade-offs between generalizability and precision. This requirement is especially acute in imaging in the life sciences, where a wide range of model organisms and imaging environments are employed ([Kamentsky et al.(2011) Kamentsky, Jones, Fraser, Bray, Logan, Madden, Ljosa, Rueden, Eliceiri, and Carpenter, Zhan et al.(2015) Zhan, Crane, Entchev, Caballero, Fernandes de Abreu, Ch’ng, and Lu, Crane et al.(2014) Crane et al.(2012), Stirman, Ou, Kurshan, Rehg, Shen, and Lu, Federici et al.(2012) Federici, Dupuy, Laplaze, Heisler, and Haseloff]). Methodologies that apply to all these diverse organisms and experimental conditions are necessarily agnostic about the constraints that are specific to a particular case. With this limitation in mind, we here confined our interest to the automated segmentation of S. cerevisiae cells in microfluidic experiments: specifically long-term imaging using devices containing individual cell traps (Fig. 1). The microfluidic device we use for this work, ALCATRAS, is from [Crane et al.(2014) Crane, Clark, Bakker, Smith, and Swain].

Budding yeast itself is a popular model organism, being a eukaryote that is easily cultivated and genetically manipulated. Widespread interest in the replicative aging of single S. cerevisiae cells has resulted in an explosion in the number of microfluidic devices that can trap mother cells for their entire lifespan ([Ryley and Pereira-Smith(2006), Sik et al.(2012)Sik, Avalos, Huberts, Lee, and Heinemann, Zhang et al.(2012) Zhang, Luo, Zou, Xie, Brandman, Ouyang, and Li, Xie et al.(2012)Xie, Zhang, Zou, Brandman, Luo, Ouyang, and Li, Crane et al.(2014) Crane, Clark, Bakker, Smith, and Swain, Jo et al.(2015)Jo, Liu, Gu, Dang, and Qin]). Budding yeast divide rapidly - growing exponentially with a doubling period of 80-90 minutes. To image the same cells over a long period of time, newborn cells (daughters) must be removed to prevent the device from becoming overcrowded. In contrast to typical cell tracking, where there is only a small probability of losing a tracked cell if it either dies or moves outside the field of view ([Li and Kanade(2007)]), this removal means that cells regularly appear, disappear, and replace each other. In addition, there is a stringent constraint on the speed required of any automated algorithm because experiments are long, with hundreds of cells imaged, often every five minutes, for days.

Here we present a comprehensive framework to segment and track budding yeast cells. By focusing on budding yeast in microfluidic traps, we can leverage prior knowledge about shape, motion and appearance to improve accuracy and performance. This approach, employing both fitted probabilistic models and supervised machine learning, is generally applicable and can provide substantial improvements in accuracy.

## Approach

Our framework is structured into four stages for integrated identification, segmentation and tracking of cells:

1. the microfluidic features of the traps are located and used to define regions of interest and to register images
2. probable cell seeds are identified using a supervised learning classifier applied to each pixel
3. the location of cells at previous time points is used to improve the predicted cell seed locations
4. a shape based active contour model is iteratively applied to proposed seeds until the image is segmented.

**Figure 1.**
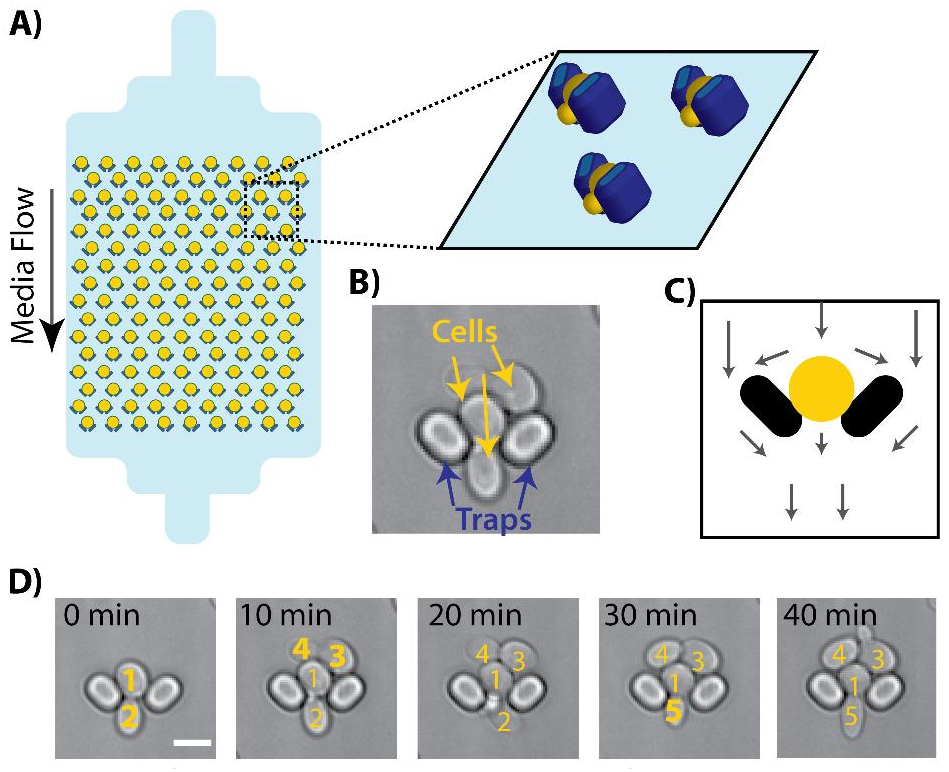
Microfluidic devices for long term imaging of cells impose physical constraints. A) A microfluidic device for budding yeast where cells are held in place by media flow and imaged over long periods of time. B) An image of a single trap containing multiple cells. The cells and traps share many similarities in shape and optical properties. C) The trap design imposes specific physical constraints on where cells can be located and where they are likely to move. Arrows represent fluid flow in the device. D)Time-lapse images of a single trap shows the appearance of new cells washed in from above, and the disappearance of daughters that are washed away after birth. Cells are individually labelled to show the continuity between time points and the appearance of new cells (bold). Scale bar is 5µ m.

## Methods

### Identifying physical features of the microfluidic device

The widely used microfluidic devices with traps have floor to ceiling pillars that hold cells and create regular optical features ([Chen et al.(2016) Chen, Crane, and Kaeberlein]). These microfluidic features are not only consistent, stable landmarks, but they constrain cell motion in a predicable manner. We therefore use these physical landmarks at all stages of processing to inform and constrain the segmentation to increase precision.

To locate the microfluidic features, the software predicts the locations of traps by performing normalized cross-correlation between the initial time point of the experiment and a canonical image of the microfluidic features. Following this prediction, user feedback is required to correct (add or remove) any features that were not accurately detected. The importance of identifying trap locations mandates input from the user, but this input is only performed at the initial time point and as such is not laborious. Following this identification, the microfluidic traps are tracked through time to correct for any motion or drift from the stage of the microscope.

### Supervised classi cation to identify cell centres

Accurately determining cell centres is an important part of the segmentation because accurate cell seeding removes the need to both eliminate aberrant seeds and fuse multiple seeds into a single cell. Previous methods for budding yeast have largely used thresholding. Most commonly, a threshold is applied to either the image itself ([Gordon et al.(2007)Gordon, Colman-Lerner, Chin, Benjamin, Yu, and Brent, Pelet et al.(2012)Pelet, Dechant, Lee, van Drogen, and Peter]) or to the accumulation array of a circular Hough transform ([Kvarnström et al.(2008b)Kvarnström, Logg, Diez, Bodvard, and Käll, Dimopoulos et al.(2014) Dimopoulos, Mayer, Rudolf, and Stelling]). Alternative approaches identify foreground objects and then separate these objects with a watershed transform ([Doncic et al.(2013)Doncic, Eser, Atay, and Skotheim]). Although such methods can work well, they rely on a single feature to determine whether locations are probable cell centres and consequently can be prone to biases and under-or over-seeding. To increase the con dence in cell seeds, we designed a predictor based on a high dimensional feature space for each pixel, a common approach in computer vision ([Grys et al.(2016)Grys, Lo, Sahin, Kraus, Morris, Boone, and Andrews]).

**Figure 2.**
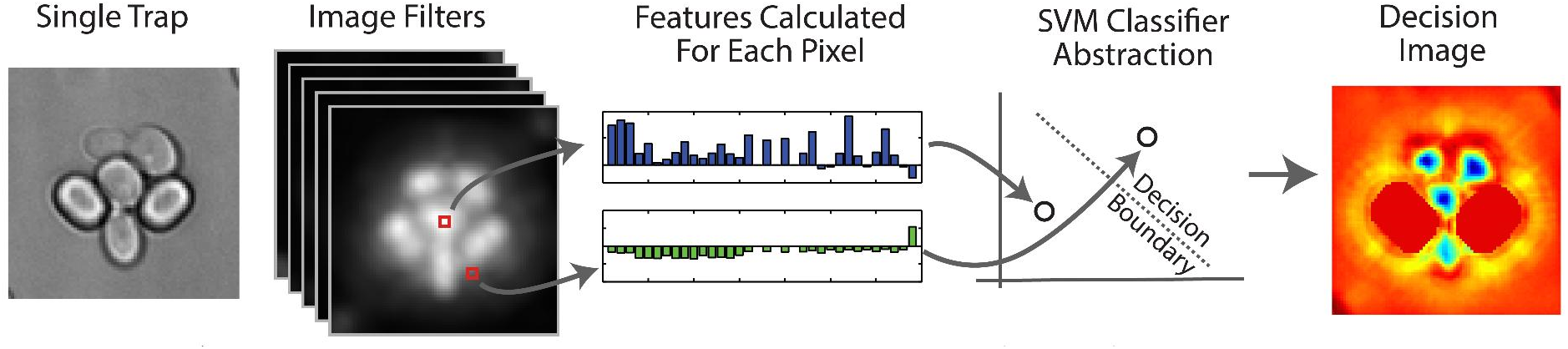
A support vector machine approach allows the use of sixty features to robustly determine probable cell centres. Two bright field images, one above and one below the plane of focus, are captured at each time point. A large range of transformations are applied to these images to generate a high dimensional feature set for each pixel. A linear SVM, trained on a curated set of images, provides a decision boundary. The signed distance of each pixel from the the decision boundary is used as a score for the cell centre. These pixel scores are reconstituted into a decision image and used to seed probable cell centres. In the image, low values (blue) indicates a better cell centreness score.

We found that the most consistent imaging conditions were generated acquiring an out of focus bright-field image ([Gordon et al.(2007)Gordon, Colman-Lerner, Chin, Benjamin, Yu, and Brent]). Although differential interference contrast (DIC) provides high contrast images, the gradient is dependent on the orientation relative to the centre of the cell which complicates segmentation ([Ning et al.(2005)Ning, Delhomme, LeCun, Piano, Bottou, and Barbano]). We acquired images both 2 μ mabove and below the central focal plane because the brightfield image is different depending on whether the image is acquired above or below the plane of focus (a bright cell with dark edges or the reverse). Both images were used in the generation of cell seeds.

To determine whether an individual pixel is likely to be the centre of a cell, transforms are run on the whole image to extract different information about the pixel and its locale. These features are then fed into a support vector machine (SVM) classifier trained on example pixels. By employing a large number of features, the classifier is better able to predict cell centres. A complete list of all sixty features used can be viewed in SOM, but include the radial Hough transform, image smoothing and sharpening features, and relational features to incorporate proximity to the microfluidic traps. Using a manually curated ground-truth data set, we trained a linear-SVM using the publicly available liblinear library ([en Fan et al.(2008)en Fan, wei Chang, jui Hsieh, rui Wang, and jen Lin]). Both polynomial and RBF-kernel SVMs (using the libSVM library) were tested, but offered negligible improvements in accuracy despite dramatically increasing run-time. To determine the decision boundary, we define λ to be a slack variable that constrains the cost of misclassification, the training set as the set of vector-label pairs 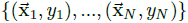, then the support vector, 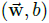, is selected to minimize

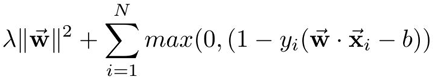

Where 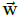 is the vector containing the weight for each image feature used in the classification and *b* is an offset. The score of each pixel is determined with the function, *g*(*x*), given by:

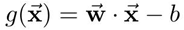

We use a five-fold cross-validation approach to determine the cost parameter, λ.

For a given image to be segmented, the features are calculated and passed to the trained SVM to generate a score for ‘cell-centreness’ for each pixel. The scores are reconstituted into an image of the same size as the original where low values indicate pixels likely to be cell centres (Fig. 2). We refer to this image as the decision image.

### Segmentation using a morphologically constrained cell-shape model

Although many cell types have a highly constrained morphology, most segmentation algorithms do not use this information to preserve generalizability ([Dimopoulos et al.(2014) Dimopoulos, Mayer, Rudolf, and Stelling]), but this choice can result in a low accuracy when imaging conditions are challenging ([Delgado-Gonzalo et al.(2015) Delgado-Gonzalo, Schmitter, Uhlmann, and Unser]). Here we employ a cell-shape model based on the morphological constraints of budding yeast, which typically have round to elliptical morphologies. Although the morphological shape space is confined for young cells, it diverges as cells age and become irregular.

Active contour methods provide a straightforward and physically motivated means of encoding shape information. They have been used extensively for image segmentation ([McInerney et al.(1996)McInerney, McInerney, Terzopoulos, and Terzopoulos, Garner(2011), Delgado-Gonzalo and Unser(2013)]), including the microscopy of *S. cerevisiae* ([Bredies and Wolinski(2011), Kvarnström et al.(2008a)Kvarnström, Logg, Diez, Bodvard, and Käll]). The boundary of a cell is defined by a deformable contour parameterised by a small number of shape parameters ([McInerney et al.(1996)McInerney, McInerney, Terzopoulos, and Terzopoulos, Blake and Isard(2012)]). The image to be segmented is processed to give a forcing image in which pixels that are likely to be part of an edge have low values. The ‘best’ contour is then found by minimising a cost function that depends on both this forcing image and the shape of the contour. If the same object is seen in multiple frames of a time-lapse, the cost function can also include terms spanning time points to punish physically improbable changes in the object’s outline. Ideally, by integrating both prior knowledge about the physiological shape space and the acquired image data, active contour methods are capable of providing increased segmentation accuracy.

We generated our forcing image from an out of focus brightfield image by taking the gradient along radial vectors from the putative centre of the newly identified cell ([Gordon et al.(2007)Gordon, Colman-Lerner, Chin, Benjamin, Yu, and Brent, Kvarnström et al.(2008b)Kvarnström, Logg, Diez, Bodvard, and Käll, Dimopoulos et al.(2014) Dimopoulos, Mayer, Rudolf, and Stelling]). This procedure highlights the edge of the cell, which appears as a white object with a black halo, but not the nearest edges of adjacent cells. To this image we add the normalised and inverted decision image so as to force the contour away from cell centres (Fig. 3B and SOM).

Given that *S. cerevisiae* cells have ovoid, concave shapes, and letting r and θ be the usual polar coordinates, *s* be a periodic cubic B spline with six evenly spaced knots, r, in the range 0 to 2π, we defined our contour as all pixels intersected by the curve (Figure 3Aii):

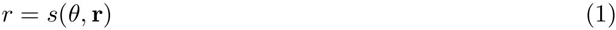

This definition allows a range of physically reasonable cell shapes with only six parameters – the six elements of the vector r – and balances the competing interests of complexity and flexibility ([Delgado-Gonzalo and Unser(2013)]).

We used a dataset of manually curated cell shapes to determine the empirical distribution of the parameters of the morphological shape space. Specifically, a multivariate normal distribution was fitted for the parameter vector r and added to the cost function for new cells to punish unphysical cell shapes. If *F* is the forcing image and 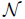 is the probability density function of the normal distribution with parameters μ and Σ fitted to the curated data, then the cost function becomes

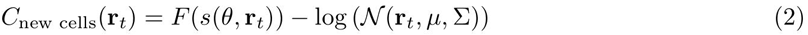

For tracked cells, we find that it is advantageous to punish unphysical changes in shape and to include cell growth. Cells are more likely to grow than shrink, although often stay the same shape. To capture these we effects we use a log-normal distribution because of its positive skewness. Fitting the distribution to the element wise division of the parameter vector for the cell at the current time point (r_t_) by the parameter vector for the same cell at the previous time point (r_t−1_) punishes the relative change in shape, rather than the absolute, which improves the outline Identification for large cells. We curated a time-lapse data set to t the multivariate log-normal distribution to the element wise division. Writing this element-wise division as 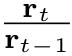 and ln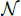 as the probability density function of the log-normal distribution with parameters *μ*^′^ and ∑^′^ fitted to the curated time-lapse data, the nal cost function is:

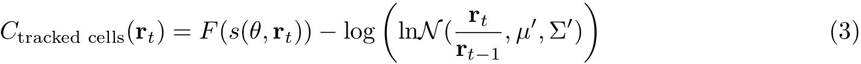

**Figure 3.**
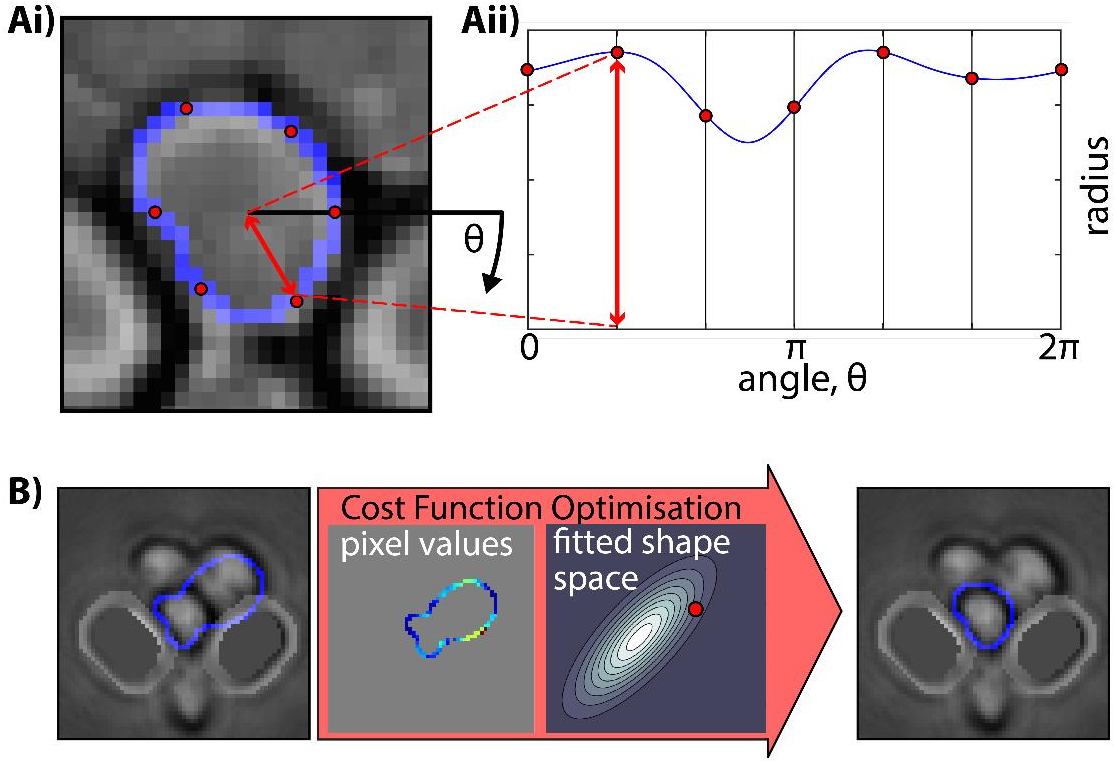
Using a morphological model for cell segmentation allows prior information about cell shape to be exploited. A) Definition of cell contour: The contour is defined by a periodic cubic B spline in polar coordinates (blue line in Aii) centred on the cell seed, and is completely defined by the six knots of the spline (red circles in Aii). This spline is mapped back to the image coordinates to produce a cell contour (Ai). B) Identification: To identify the correct contour for a particular cell a forcing image is generated by taking the gradient of the image in the radial direction from the cell seed (B, left). The contour is found by optimising a cost function that combines the value of the pixels in the forcing image along the cell contour with the probability of the contour according to a fitted probability distribution of cell shapes.

The JarqueBera test ([Jarque and Bera(1980)]) was applied to data fitted to a normal distribution to ensure the distribution appropriately modelled the data; details can be found in the SOM.

The boundary of the cell is found by globally optimising this cost function for r_t_ using a particle swarm ([Birge(2003)], Fig. 3B).

### Incorporating temporal information to refine cell centre prediction

Budding yeast are constantly growing and dividing, and coupling temporal tracking information with knowledge about fluid flow can increase the accuracy of cell Identification. Fluid flow on the small length scales of microfluidic devices has a low Reynolds number and so is predictable and consistent. The cell traps and predictable flow profile affects both where cells are initially located and where cells are likely to move to as they grow.

For time points after the first, we developed a method that incorporates this prior knowledge about the physical imaging platform. We generate a prior image *m(x, y)* for the motion of each cell at the previous time point, which encodes the probability that the the centre of cell has moved to the point *(x, y)* at the current time point.

**Figure 4.**
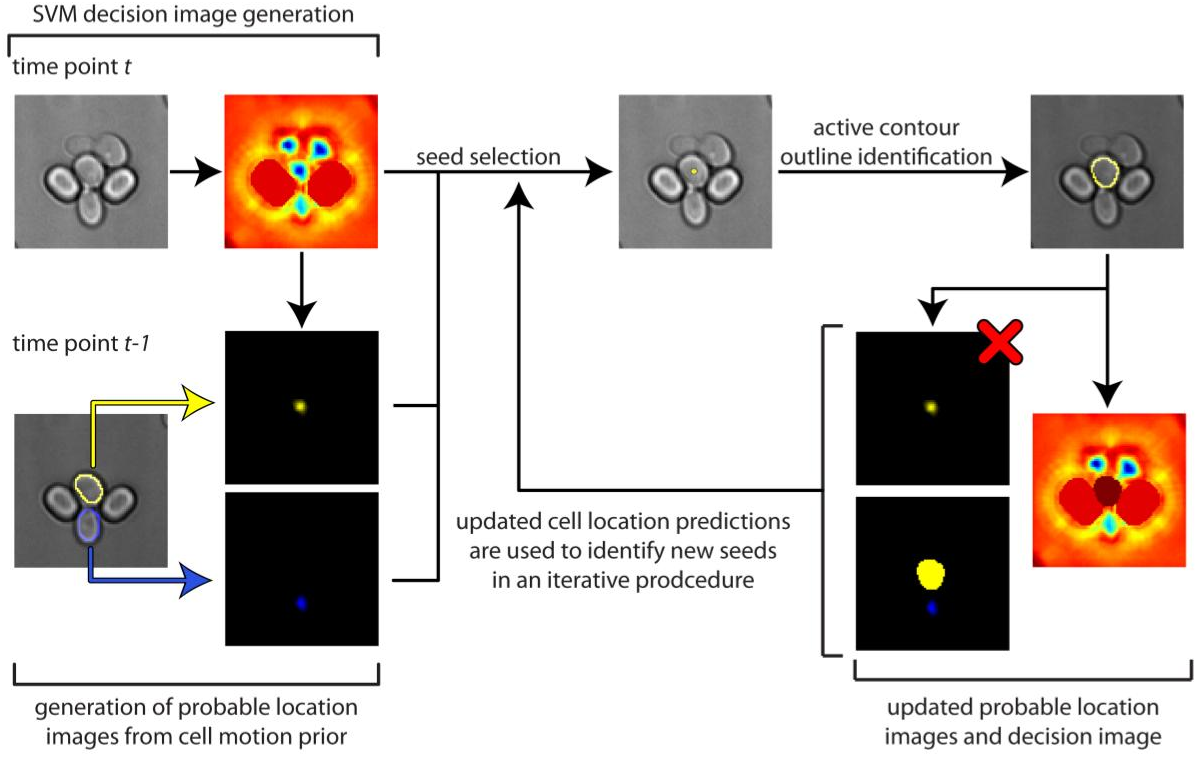
Greedy optimisation prioritises cells identified at previous time points to ensure reliable tracking. To segment cells at current time point (*t*), a decision image and probable location images are generated as described in the main text and subjected to a greedy optimisation algorithm. The probable location images are used to propose a seed: the highest scoring pixel in the image set. If it is above a threshold value, the seed is used to generate a cell contour as described in Figure 3. If the contour meets criteria for its shape and its overall score, it is stored, and the decision and probable location images updated to prevent new cells being found in the region it occupies. If it is a tracked cell, the probable location image for this cell is no longer considered during seed generation. The procedure is repeated until no pixels above the threshold criteria remain. From this point on *new* cell seeds are proposed from the decision image.

The motion prior is indexed by a cell’s size and location in the trap and is generated from empirical measurements: the curated pairs of cells used to train the distribution of tracked cell shapes (equation 3). When a motion prior is calculated for a particular cell, two probability density functions are retrieved – one indexed by its size and one indexed by its location – and the average returned as the motion prior for that particular cell.

To combine this motion prior with the likely cell locations at the current time point, a probable location image is calculated for each cell as:

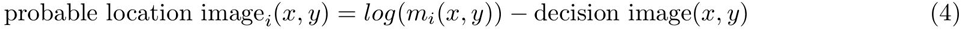

An example set of probable location images is shown in figure 4.

### Iterative greedy optimization of cell contours

To try to maximize the performance of the segmentation, we use a greedy optimization that integrates information about cell centres (decision image) and cell shape (active contour algorithm). For the first time point, the decision image is calculated and the first seed selected as the pixel in the decision image with the lowest value (i.e. the pixel most likely to be a cell centre). Provided the value of this pixel is below a user-defined ‘new cell Identification’ threshold, the active contour algorithm is applied using the cost function given by equation 2 and returns the outline of the putative cell and its score: the value of the cost function. If the cost is below a user-defined threshold, the cell is deemed to be a true cell and assigned a unique label for tracking. The cell is then blotted out of the decision image so that no new cell seeds will be identified within the area of the previously identified cells. The procedure is repeated until no pixels remain that are below the user-defined threshold.

At subsequent time points, the set of probable location images is used to generate cell seeds. A similar greedy optimisation is applied to these images: identifying the highest scoring pixel in the image set, applying the active contour algorithm with the tracked cell cost function (equation 3), and storing the cell if its score and change in shape meet appropriate threshold criteria. With each successful Identification, the probable location and decision images are modified to include the newly identified cell. Once all previously identified cells have been tracked, new cells are identified by applying the first time point iterative procedure to the modified decision image.

This division into tracked cells and new cells has a number of advantages:

1. it improves consistency in the location and shape of the cells across time by using shape and location information of cells at previous time points to identify and segment cells at the current time point
2. it reduces false positives in new cells and false negatives for cells present over multiple time point by allowing both a more lenient criteria to be applied to cells identified at the previous time point and a more stringent one to be applied to new cells
3. it helps prevent large and irregularly shaped older mothers from being confused for multiple smaller new cells by allowing us to delineate the new-cell and existing cell shape models.

This procedure is applied iteratively over all time points, segmenting the time-lapse images and tracking the cells. The algorithm is shown schematically in figure 4; further details, with pseudo code, can be found in the SOM.

### Methods comparison

To evaluate the performance of our framework, we generated and manually curated several data sets for shape and tracking of individual cells. To maintain independence and prevent biasing of the measures of performance, we used strains in which a fluorescent reporter was strongly expressed in the cytoplasm. This fluorescent reporter was used for segmentation by applying a circular Hough transform and Chan-Vese active contour algorithm: reducing the overall workload and ensuring the initial segmentation was independent of any one method to be tested. Following segmentation of the fluorescent channel, we manually curated all outlines using brightfield images. To ensure the datasets were representative of the variability seen in experiments, images were acquired over multiple months in different conditions and by different individuals (see SOM). These datasets are separate from those used for training DISCO. The curation was done in two parts: one for cell shape and size error rates and a second for tracking error rates between time points. The final data sets used for error determination contained >1,000 manually curated cell outlines and >1,200 curated cell trajectories.

The performance statistics for segmentation and tracking were taken from the ISBI cell tracking challenge ([Maška et al.(2014)Maška, Ulman, Svoboda, Matula, Matula, Ederra, Urbiola, Españna, Venkate-san, Balak, Karas, Bolcková, Streitová, Carthel, Coraluppi, Harder, Rohr, Magnusson, Jaldén, Blau, Dzyubachyk, Kížek, Hagen, Pastor-Escuredo, Jimenez-Carretero, Ledesma-Carbayo, Muñoz-Barrutia, Meijering, Kozubek, and Ortiz-de Solorzano]). These performance measurements result in a single comprehensive score for tracking and segmentation, which makes it possible to compare methods directly. The metric for segmentation accuracy is the Jaccard index and is given by:

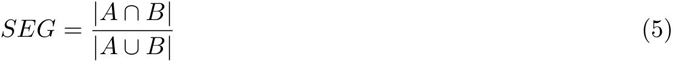

Where *A* is the ground truth segmentation for that cell and *B* is the cell outline identified by the test algorithm. The inclusion of the intersect and union means that this score punishes over-segmentation and under-segmentation to the same degree. Cells that are either false positive or false negatives receive a score of 0; cells that are perfectly segmented receive a score of 1.

For estimating the accuracy of tracking, the measurement is based on transformations applied to an acyclic oriented graph ([Maška et al.(2014)Maška, Ulman, Svoboda, Matula, Matula, Ederra, Urbiola, Españna, Venkatesan, Balak, Karas, Bolcková, Streitová, Carthel, Coraluppi, Harder, Rohr, Magnusson, Jaldén, Blau, Dzyubachyk, Kížek, Hagen, Pastor-Escuredo, Jimenez-Carretero, Ledesma-Carbayo, Muñoz-Barrutia, Meijering, Kozubek, and Ortiz-de Solorzano, Matula et al.(2015)Matula, Maška, Sorokin, Matula, Ortiz-de Solórzano, and Kozubek]). Each node in the graph is a detected cell at a specific time point, and the edges connect cells identified to be the same at different time points. The error for an individual cell is determined by the number of operations that must be performed to make the acyclic oriented graph for the test data set match the ground truth. Operations are weighted by the time required to perform them manually (i.e. more mouse clicks incur a higher cost). The tracking cost for each individual cell is normalized by the number of time points for which it is present to give a score between 0 and 1.

**Figure 5.**
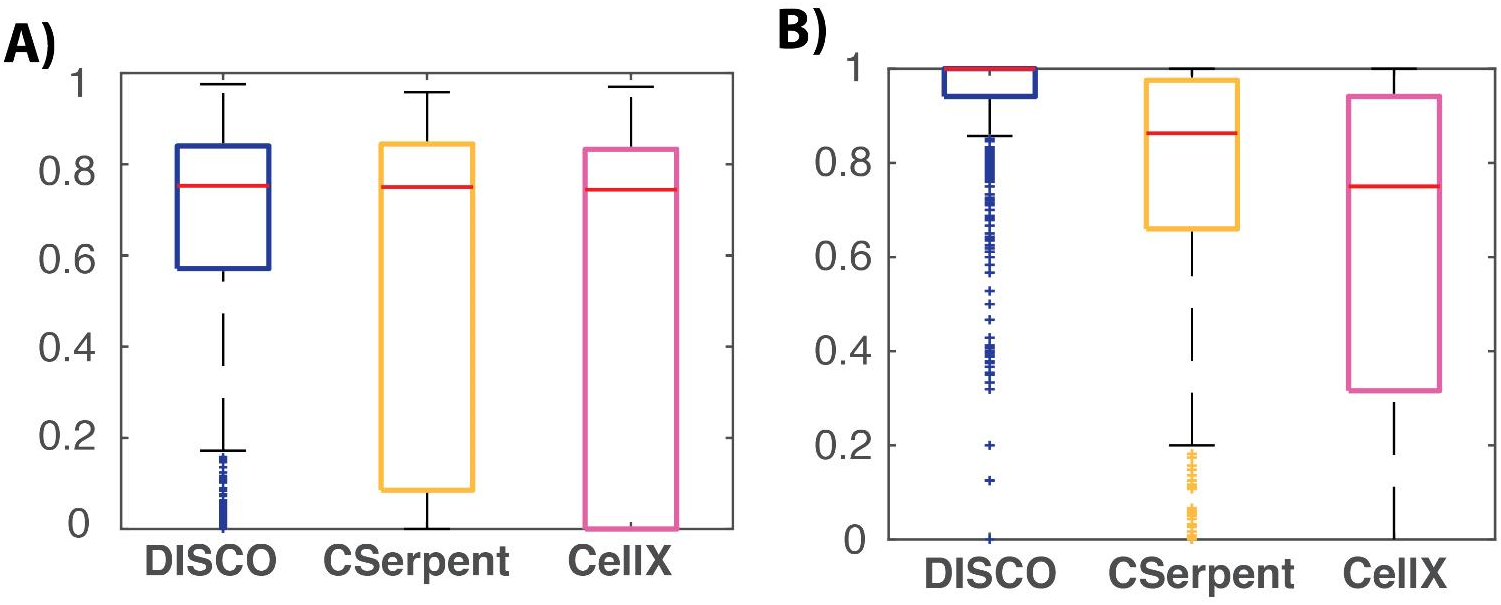
Segmentation and tracking performance comparison for DISCO, CellX and CellSerpent. A) Segmentation accuracy comparisons. Each of the three software packages were analyzed on all cells in the ground-truth data set and the Jacc-card index was used for scoring. B) Tracking accuracy comparisons. DISCO and CellX were run using the native tracking algorithms, and CellSerpent was run using an overlap tracking approach. The significant performance differences illustrates the confusing nature of these images to current segmentation approaches.

For comparison, we selected the CellX ([Dimopoulos et al.(2014) Dimopoulos, Mayer, Rudolf, and Stelling]) and CellSerpent ([Bredies and Wolinski(2011)]) software. Although CellSerpent has a similar structure to our algorithm with the proposal of cell seeds followed by an active contour procedure to identify cell boundaries, it is different in that a single heuristic feature (the circular Hough transform) is used to identify seeds, no time information is employed, cell shape is enforced simply by punishing deviations from circularity, and the contour is optimised locally, not globally, to identify the cell outline. CellX, in contrast, is substantially different from our software. Although CellX uses a Hough-based seeding too, the cell outline is found by a graph cut algorithm applied to an edge image generated using a membrane profile and the proposed seed. This approach imposes little shape constraint on the cell. Both CellX and CellSerpent were designed for single layers of cells, tightly confined in the vertical axis. For comparing tracking, we modified CellSerpent to enable a common tracking methodology because tracking is not enabled by default. We use the overlap between cells at adjacent time points and assign the same cell labels if the overlap is > 0.5. For CellX, the tracking provided with their GUI was used. Prior to comparison, we attempted to optimize the performance of both packages on each of the test data sets according to the instructions provided and, for CellX, with input from the authors. To ensure fair comparison, the results of the CellX and CellSerpent segmentation for each experiment were uniformly dilated or eroded over a range of sizes and scored. We use the best performing segmentation score for each test data set in the comparison.

Both alternatives perform worse than our approach (Fig. 5). The performance of the three algorithms illustrates how different the images from the microfluidic system are from those of single layers of cells for which CellX and CellSerpent were designed. The slight freedom in z motion makes a consistent membrane profile difficult to define, complicating detecting cell centres and increasing the importance of the time and morphology information we incorporate. In addition, both alternatives were slightly slower than DISCO (SOM).

## Discussion

Image segmentation and tracking has been widely recognized as a pressing problem that must be addressed to enable high-throughput, high content image cytometry. Many previous approaches assume a uniform field of view and fail to incorporate prior information. With the increasing popularity of microfluidic methods, there is a need for approaches that are able to use the a priori information of physical constraints imposed by the microfluidic systems to improve segmentation and tracking. We believe that this work, which combines aspects of supervised classification with model driven segmentation, contributes to this aim.

Moving from a single-feature threshold for determining cell centres and seeds to a multi-feature supervised classifier offers several advantages. Although single-feature heuristic approaches are appealingly simple, they result in substantially lower performance and require a high degree of user intervention to optimize performance for separate experiments. Furthermore, by increasing the accuracy of predicting cell seeds, we were able to rely more heavily on image features during segmentation and better exploit shape information. Taking advantage of this information can provide significant improvements (Fig. 5). Rather than relying solely on optical features, we employ the substantial prior knowledge available about our imaging system and biological samples to increase performance. The inclusion of temporal and morphological models provides reliable segmentation and tracking, even in difficult conditions. While shape and model based segmentation approaches are widely used, our algorithm fits generative models to empirically curated datasets: improving robustness and reducing the need for parameter tuning. Given that they are fitted to curated data sets, we expect that the same methods can be straightforwardly applied to other devices and experimental setups.

Microfluidic devices have the potential to provide unprecedented quantities of high content data, especially for aging research: an area that been low throughput because of the effort in performing manual micro-dissection ([Kaeberlein(2010)]). Nevertheless, the widespread adoption of these systems will depend on having the capability to robustly and consistently process the information in the acquired images, using frameworks such as the one we present here.

## Acknowledgement

We would like to acknowledge our funders for providing us with the means to pursue this work, Swain lab members for constructive comments on the manuscript and Lucia Bandiera for providing some of the images used in the performance comparison.

This work was supported by the BBSRC & HFSP (MMC, EB & PSS) & SULSA (EB)

